# New loci and neuronal pathways for resilience to heat stress in animals

**DOI:** 10.1101/2021.02.04.429719

**Authors:** Evans K. Cheruiyot, Mekonnen Haile-Mariam, Benjamin G. Cocks, Iona M. MacLeod, Ruidong Xiang, Jennie E. Pryce

## Abstract

Climate change and resilience to warming climates have implications for humans, livestock, and wildlife. The genetic mechanisms that confer thermotolerance to mammals are still not well characterized. We used dairy cows as a model to study heat tolerance because they are lactating, and therefore often prone to thermal stress. The data comprised almost 0.5 million milk records (milk, fat, and proteins) of 29,107 Australian Holsteins, each having around 15 million imputed sequence variants. Dairy animals often reduce their milk production when temperature and humidity rise; thus, the phenotypes used to measure an individual’s heat tolerance were defined as the rate of milk production decline (slope traits) with a rising temperature-humidity index. With these slope traits, we performed a genome-wide association study (GWAS) using different approaches, including conditional analyses, to correct for the relationship between heat tolerance and level of milk production. The results revealed multiple novel loci for heat tolerance, including 61 potential functional variants at sites highly conserved across vertebrate species. Moreover, it was interesting that specific candidate variants and genes are related to the neuronal system (*ITPR1, ITPR2,* and *GRIA4*) and neuroactive ligand-receptor interaction functions for heat tolerance (*NPFFR2, CALCR,* and *GHR*), providing a novel insight that can help to develop genetic and management approaches to combat heat stress.

**Author summary:** While understanding the genetic basis of heat tolerance is crucial in the context of global warming’s effect on humans, livestock, and wildlife, the specific genetic variants and biological features that confer thermotolerance in animals are still not well characterized. The ability to tolerate heat varies across individuals, with substantial genetic control of this complex trait. Dairy cattle are excellent model in which to find genes associated with individual variations in heat tolerance since they significantly suffer from heat stress due to the metabolic heat of lactation. By genome-wide association studies of more than 29,000 cows with 15 million sequence variants and controlled phenotype measurements, we identify many new loci associated with heat tolerance. The biological functions of these loci are linked to the neuronal system and neuroactive ligand-receptor interaction functions. Also, several putative causal mutations for heat tolerance are at genomic sites that are otherwise evolutionarily conserved across 100 vertebrate species. Overall, our findings provide new insight into the molecular and biological basis of heat tolerance that can help to develop genetic and management approaches to combat heat stress.

## Introduction

Heat stress from rising global temperatures is an issue of growing importance across tropical and temperate zones affecting humans, livestock, wildlife, and plants. A recent study [1] indicates that many people are now exposed to harmful heat, and this has risen by more than two-fold when compared to the pre-industrial climates (i.e., 95 versus 275 million people), with future projections showing that over 1 billion people will experience an even greater impact of heat within the next 50 years [2]. In livestock, the annual temperature-humidity values that rise above thresholds considered to be comfortable have been increasing in many regions including Australia, the USA, Canada, and parts of Europe [3, 4], making heat stress a multimillion-dollar issue in the livestock industry that compromises production (reduced growth, milk, eggs, etc.) and reproduction leading to economic losses [5].

The thermoregulatory capacities of mammals and plants to cope with extreme heat have been studied for decades. Genetic variation of thermoregulation during heat stress exists within species, including cattle breeds, with the literature indicating that tropical breeds, such as Zebu (*Bos indicus*), have a better tolerance to temperature and humidity than cattle from temperate zones (e.g., Holsteins), in part, due to the lower productivity of Zebu cattle [6]. Temperate breeds also show genetic variation in heat tolerance; for example, New Zealand Holsteins appear to exhibit higher reductions in milk yields in hotter climates than Jerseys or crossbreds [7]. While it is not fully understood why animals differ in their thermotolerance, it is hypothesised to be due to a myriad of biological mechanisms; including cellular, morphological (coat color, coat length, etc.), behavioural (e.g., feed and water intake, standing and lying time), as well as neuro-endocrine systems. See comprehensive review by [8] for more information. Notably, the molecular basis for differences in these adaptive responses within various mammalian species is still largely unknown.

Dairy cattle are excellent and convenient model for enhancing our knowledge on the molecular aspects of heat tolerance in mammals for two main reasons: 1) large phenotype datasets needed to study heat tolerance, as well as extensive genomic information, are available; 2) they have been genetically selected mainly for high milk production over many years, offering an opportunity to understand the genetic basis for coping with both environmental and elevated metabolic-heat stress associated with increased milk production.

The development of methods to describe heat tolerance in cattle has been an active research area for many years. Measuring changes in core body temperature (e.g., rectal, vaginal, rumen temperature, etc.), thermal indices (e.g., temperature-humidity index (**THI**)) are some of the ways to assess thermal adaptations and performance in animals. [9] pioneered using daily milk yield and temperature-humidity data to measure variability in the rate of decline in milk yield associated with variability in response to heat stress. This method has been widely adopted due to the availability of large datasets from routine recording in dairy farms, e.g., [3]. Heat tolerance in dairy cattle measured using rectal temperatures or the rate of milk yield decline is partly under genetic control, having a low (0.1) to moderate heritability (0.30) [3, 9, 10], which makes it amenable to selection. As such, considerable research has been undertaken to provide breeding solutions for heat stress, which is already a feature of dairy cattle breeding programmes in some parts of the world, e.g., Australia [3]. Identifying specific genetic variants that increase tolerance to heat may help to improve dairy breeding programmes in addition to improving our knowledge of the thermal biology in other mammals. However, except for mutations in the SLICK locus [11], the identification of the specific genetic variants for heat tolerance in cattle and other species has, in most cases, remained elusive, in part due to many reasons, including the sample size used in past studies.

Having a large sample size is particularly important for identifying rare causal variants with medium-sized effects and common variants with small effects. As sample size increases, the loci significantly associated with complex traits are expected to increase, as demonstrated for the human height [12]. Several genome-wide association studies (**GWAS**) using Single nucleotide polymorphisms (**SNPs**) have been conducted over the last decade to identify candidate causal genes for various heat tolerance traits (rectal temperature, heart rate, sweating rate, rate of milk yield decline, etc.) in dairy cattle [13–16] and pigs [17]. However, these GWAS were underpowered, with the largest sample size to date of around 5,000 animals [13, 14]. These studies have also used standard industry SNP panels of random genome-wide markers, either 50k or 600k SNPs, leading to inconsistencies and poor replication of the results. Although these studies have identified over 400 significant variants associated with heat stress in animals, none were established to be causal mutations.

Here, we performed a GWAS using milk production records of 29,107 Holstein cows, each having over 15 million sequence variants that were imputed from various lower density SNP chips to whole-genome sequence using a reference dataset of sequences from the Run7 of 1000 Bull Genome Project [18]. The specific aims of the study were to 1) perform single-trait GWAS to identify genomic variants associated with sensitivity of milk traits (milk, protein, and fat) to heat stress 2) combine single-trait GWAS results in a multi-trait meta-analysis to boost the power and identify pleiotropic variants associated with all the milk traits 3) conduct post-GWAS pathway analysis using the list of candidate genes identified in single-trait GWAS and meta-analysis to elucidate biological mechanisms underlying heat tolerance.

## Results

### Descriptive statistics and genomic heritability of the study phenotypes

The average milk, fat, and protein yields used to derive heat tolerance proxy-phenotypes (i.e., slope traits) and intercepts (representing level of milk production) are in Table 1. The slope traits derived from the milk, fat, and proteins yields using reaction norm models on a function of the temperature-humidity index (**THI**) were defined as follows: heat tolerance milk (**HTMYslope**), fat (**HTFYslope**), and protein (**HTPYslope**) yield slope traits, respectively. On the other hand, the intercept solutions from the reaction norm models – representing the level of milk production were defined as milk (**MYint**), fat (**FYint**), and protein (**PYint**) yield intercept traits. The genomic heritability estimates for the intercept traits were high [0.36 ± 0.01 (MYint), 0.30 ± 0.01 (FYint), 0.24 ± 0.01 (PYint)] compared to slope traits [0.23 ± 0.01 (HTMYslope), 0.21 ± 0.01 (HTFYslope), 0.20 ± 0.01 (HTPYslope)] (Table 1). The phenotypic correlations between the intercept and slope traits were high, with values of −0.71 (MYint versus HTMYslope), −0.77 (FYint versus HTFYslope), and −0.83 (PYint versus HTPYslope), suggesting that lower producing cows have a smaller reduction in their yield as the THI increases. The Pearson correlations of slope solutions from the reaction norm model were 0.90 (HTMYslope versus HTPYslope), 0.56 (HTMYslope versus HTFYslope) and 0.62 (HTPYslope versus HTFYslope).

**Table 1.** Average daily milk (liters), fat (kg), and protein (kg) yield for 29,107 cows and the mean of their corresponding intercept and slope traits derived from reaction norm models. The additive genetic variance (AG) and genomic heritability (h^2^) were estimated for cows based on 50k SNP panel.

|  |  | Milk yield intercept traits<sup>1</sup> |  |  |  | Milk yield slope traits<sup>2</sup> |  |  |
| --- | --- | --- | --- | --- | --- | --- | --- | --- |
| Trait | | Raw mean ± | Mean* ± | AG ± SE | $h^2 \pm$ SE | Mean* ± | AG ± | $h^2 \pm$ SE |
|  |  | SD | SD |  |  | SD | SE |  |
| Milk yield (liters) |  | 25.85 (8.19) | 26.2 (22.29) | 381.0 (14.36) | 0.36 (0.01) | 3.43 (2.92) | 4.80 (0.20) | 0.23 (0.01) |
| Fat yield (kg) |  | 0.98 (0.30) | 8.72 (7.25) | 38.78 (1.64) | 0.30 (0.01) | 1.24 (1.04) | 0.56 (0.02) | 0.21 (0.01) |
| Protein yield (kg) |  | 0.85 (0.26) | 6.96 (5.91) | 18.60 (0.89) | 0.24 (0.01) | 1.18 (0.99) | 0.47 (0.02) | 0.20 (0.01) |

\*Means were calculated based on the absolute values of each intercept and slope traits; <sup>1</sup>represents the level of milk production of cows; <sup>2</sup>heat tolerance proxy-phenotypes.

### Single-trait GWAS for intercept and slope traits

The number of significant SNPs was generally more for intercept than slope traits at the p-value thresholds tested (Table 2). At a stringent p-value of < 1E-05, the false discovery rate (**FDR**) varied between 0.02 and 0.03 for intercept and 0.02 and 0.05 for slope traits. The number of significant independent QTL (based on the number of 5 Mb non-overlapping windows across the chromosome with at least one significant SNP) ranged from 28 to 72 for intercept traits and from 21 to 37 for slope traits. At a relaxed cut-off threshold, where the FDR was < 0.10, the number of significant QTLs from single-trait GWAS ranged from 78 to 188 (intercept traits) and from 51 to 109 (slope traits).

**Table 2.**
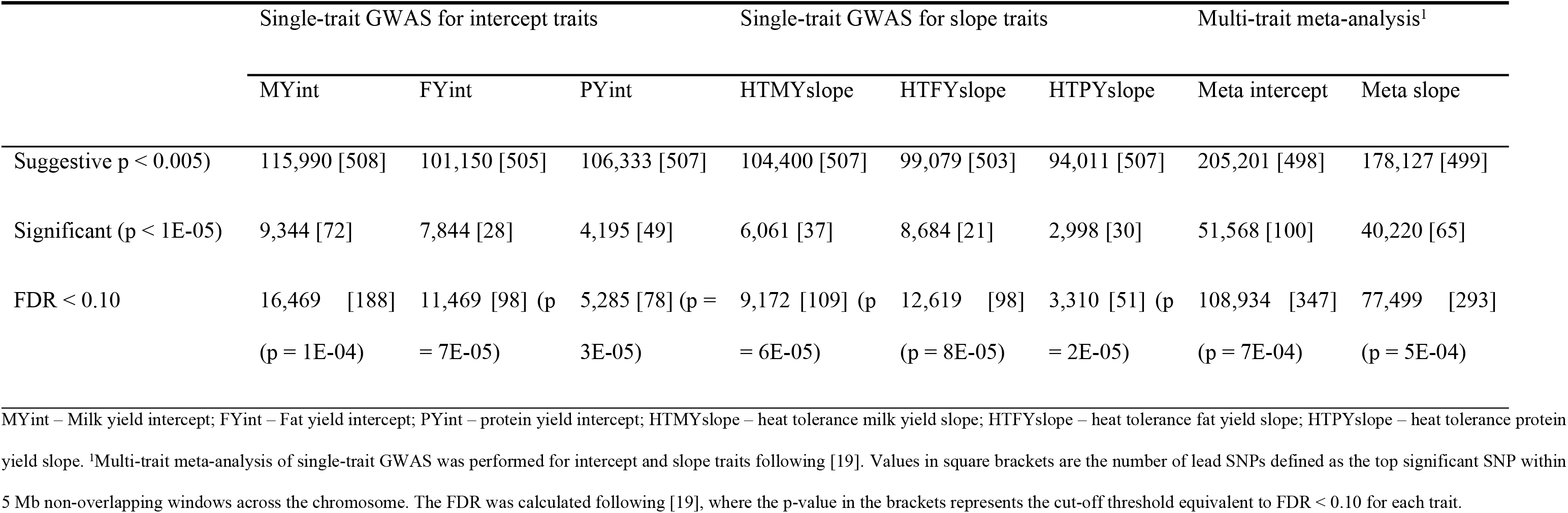
Number of SNPs identified at a p-value of < 0.005 (considered suggestive), and a p-value of < 1E-05 (significant), and false discovery rate (FDR < 0.10) for QTL discovery cows (N = 29,107) based on 15 million imputed-whole genome sequence variants.

The number of significant (p < 1E-05) QTLs (i.e., 5 Mb windows) varied across the three slope traits with greater overlap between HTMYslope and HTPYslope (13 QTLs; 20.6%) compared to HTMYslope and HTFYslope (3 QTLs; 4.8%) (S3 Fig). The overlaps were based on whether the lead SNPs (most significant) within QTLs between traits were close (within 1 Mb). Surprisingly, none of the candidate QTLs overlapped between HTFYslope and HTPYslope. The effects of the lead SNPs within QTLs that overlapped between HTMYslope and HTPYslope were generally in the same direction.

### Multi-trait meta-analysis of GWAS to detect variants with pleiotropic effects

Compared to single-trait GWAS, the number of significant independent QTLs (based on 5 Mb windows with at least one significant SNP) was much higher for a multi-trait meta-analysis (Fig 1 and Table 2). At FDR < 0.10, the number of significant independent QTLs from multi-trait meta-analysis was 347 and 293 for intercept and slope traits, respectively (Table 2). At p < 1E-05, the number of significant QTLs was 100 (meta-analysis of intercept traits) and 65 (meta-analysis of slope traits). Of the significant QTLs (p < 1E-05; N = 65) for meta-analysis of slope traits, 35% (N = 23) overlapped with the candidate QTLs for single-trait GWAS analysis based on whether the lead SNP (most significant) within overlapping QTLs were close (within 1 Mb).

**Fig 1.**
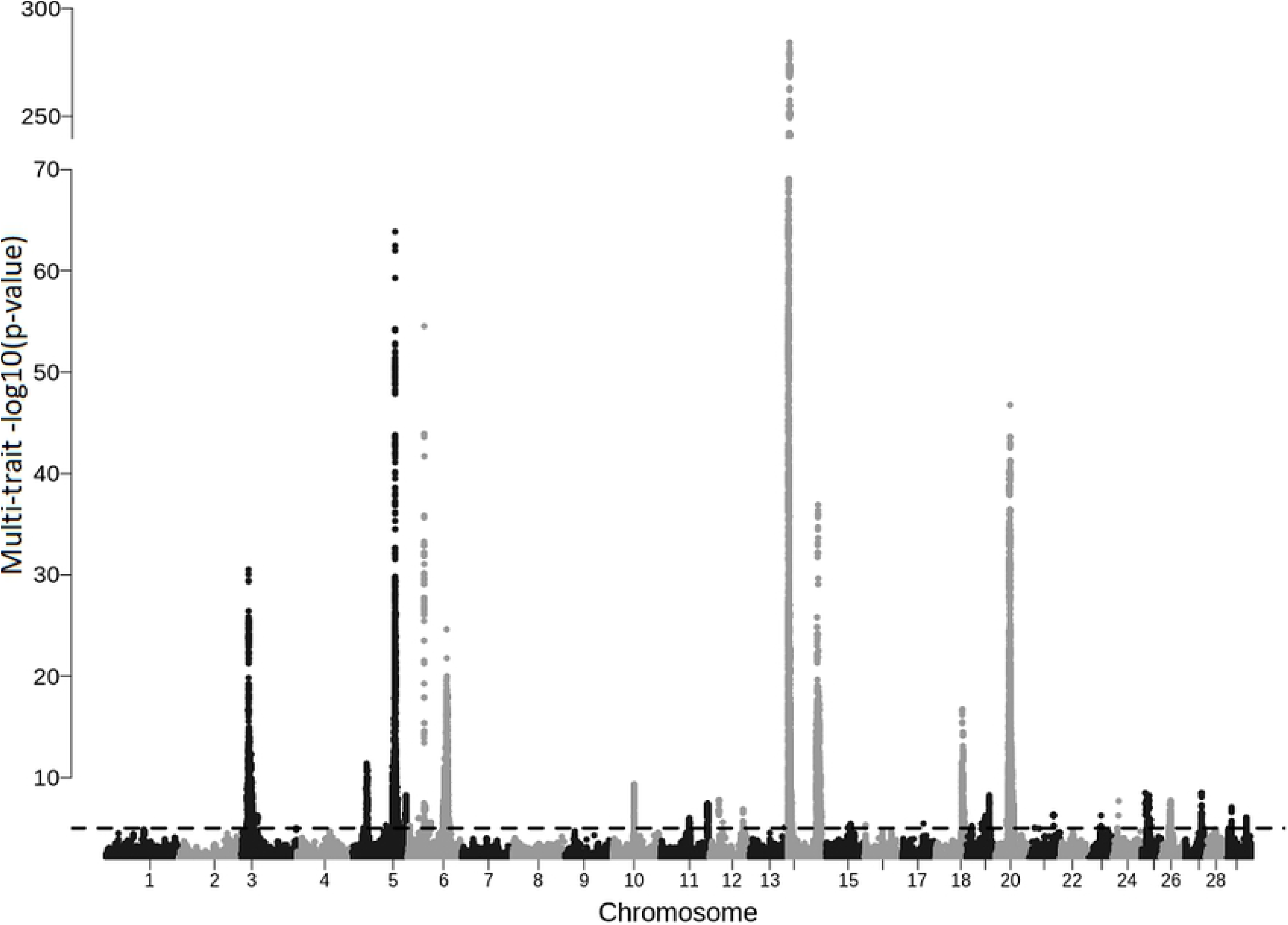
Manhattan plot of p-values obtained from combining single-trait GWAS results for milk yield slope traits.

### Lead SNPs detected using single-trait GWAS and meta-analysis of slope traits

The lead SNPs were defined as the most significant SNPs within an independent QTL (i.e., the most significant SNP chosen within 5 Mb windows across the chromosome). Detailed annotation of all the lead SNPs for single slope traits and the meta-analysis (N = 118) detected at the most stringent p-value cut-off (p < 1E-05) are in the S2 Table.

About half the lead SNPs (51%) for slopes were in relatively low LD (r^2^ < 0.5) with nearby (within 1 Mb region) lead SNPs for intercepts, indicating that they are not strongly associated with the level of milk production. Some lead SNPs mapped within or close to several candidate genes, which have been linked to environmental stress or heat tolerance in animals in previous studies, including *REG3A* [20], *NPFFR2* [21], and *CLSTN2* [22]. Several other lead SNPs mapped close to novel candidate genes that, to our knowledge, have not been described for thermotolerance in previous studies.

However, the remaining lead SNPs (49%) for slopes were in medium to strong LD (r^2^ > 0.50) with nearby (within 1 Mb) lead SNPs identified for intercept traits (S4 Fig), suggesting that they affect both traits, which was expected due to the strong genetic negative correlation between heat tolerance and milk production, with estimates in this study of around −0.80. The most significant lead SNPs for heat tolerance (slope traits) that were strongly (LD; r^2^ > 0.8) associated with the level of milk production (intercept traits) mapped close to or are within genomic loci previously reported to have pleiotropic effects on bovine milk production traits, including the *DGAT1* [23, 24], *MGST1* [25], and *GHR* gene [26].

### Conditional GWAS for slope traits on either the lead SNPs or the intercept traits

We performed two conditional GWAS for slope traits to confirm whether the top hits (lead SNPs) detected in the first-round of GWAS for the slope traits were in fact discoveries of heat tolerance rather than indicators of milk yield (as the intercept and slope traits are genetically correlated). Of interest was the conditional GWAS analysis on chromosome 14, since the highly significant QTL around 0.5 Mb harbours the DGAT1 gene and the HSF1 (heat shock factor 1) gene, for which the latter has been linked to thermotolerance in Holstein cattle in different countries, including Australia [14], and the USA [15]. Notably, the lead SNPs from the first-round of GWAS for HTMYslope and HTFYslope (Chr14:581569) and HTPYslope (Chr14:555701) traits were upstream to *SLC52A2* and a synonymous variant in the *CPSF1* genes, respectively.

Fig 2 shows conditional GWAS results for chromosome 14 (around the region which showed the strongest signal in the first-round of GWAS for the slope traits – here, the conditional analyses were for slope traits on either the lead SNP or the intercept trait. In both approaches, we found that most of the SNPs were no longer significant after conditional analysis. This was the case for HTMYslope and HTPYslope slope traits, suggesting that these SNPs were possibly tagging the lead SNPs for slope traits. The lead SNP was in strong LD (r2 > 0.8) with several other variants around this QTL spanning over 10 genes (Fig 2), including variants in the HSF1 (heat shock factor 1) gene, which implies that any variant (s) around this region are possible causal mutations for heat tolerance. Nonetheless, the complex LD within this QTL region makes it difficult to pinpoint a putative causal variant (s) for heat tolerance.

**Fig 2.**
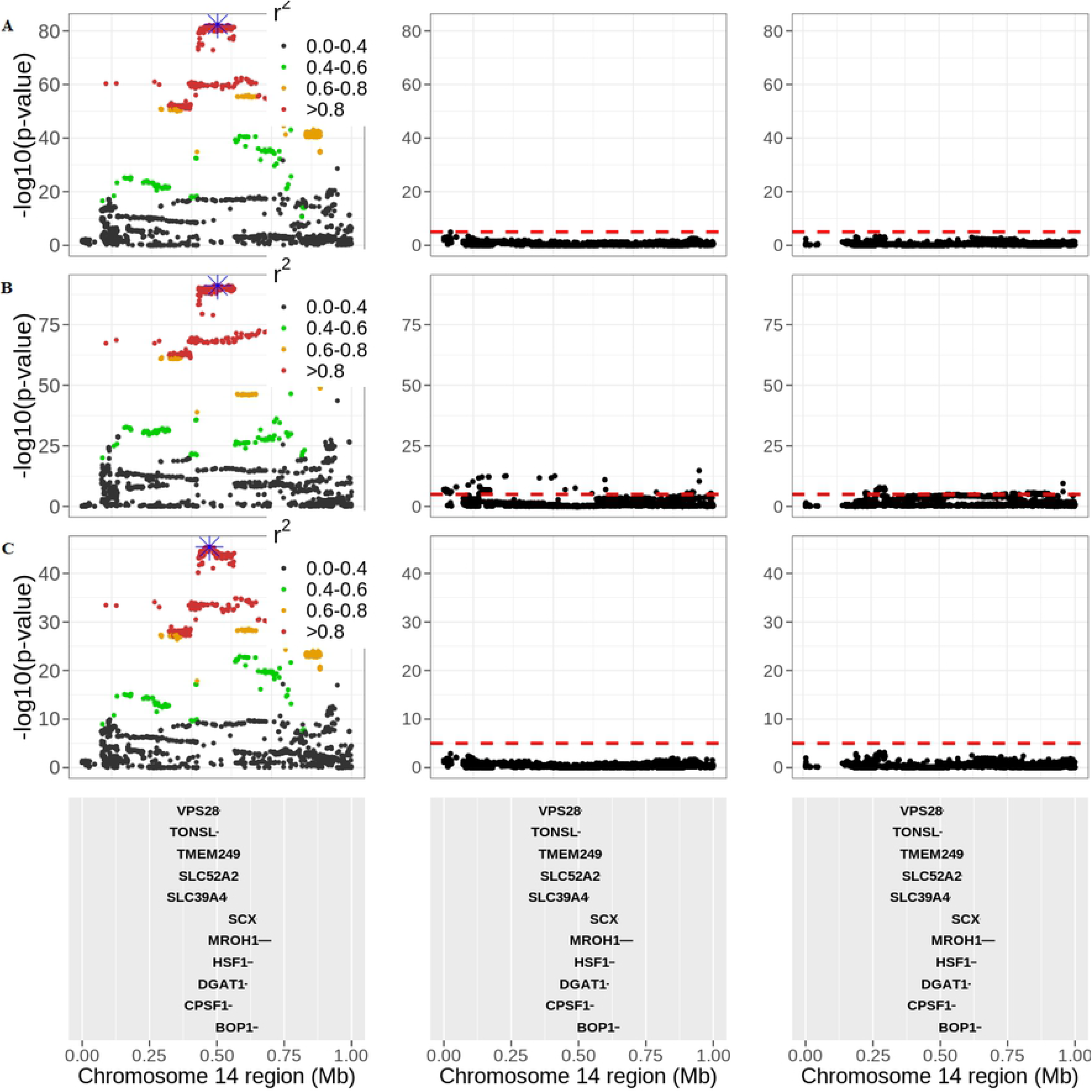
QTL discovery on chromosome 14 at 0 to 1 Mb for heat tolerance milk (HTMYslope; A), fat (HTFYslope; B), and protein (HTPYslope; C) yield slope traits. The three panels represent the GWAS p-values before conditional analysis (right panel), after conditioning slope traits on the lead SNP (highlighted in blue) defined as the most significant SNP (middle panel), and after conditioning slope traits on the intercept traits (left panel), respectively. The red horizontal dashed line is the GWAS cut-off of p < 1E-05. The strength of LD (r^2^) between the lead SNP (blue color) and all the other SNPs are color-coded accordingly.

Notably, even after fitting the lead SNP in a conditional GWAS analysis, there were still other somewhat significant (p < 1E-05) SNPs remaining for the HTFYslope trait (though not very strong signals; Fig 2), suggesting that they could be other QTLs for heat tolerance, which were not captured by the lead SNPs identified in the study.

Although the two conditional GWAS strategies (i.e., conditioning slopes on either lead SNP or intercept traits) were generally comparable regarding the strength of the GWAS signals (Fig 2), we observed a significant (Student’s t-test; p < 0.001) difference in the distribution of the GWAS p-values across slope traits. This is, in part, due to the difference in the two conditional GWAS approaches regarding the covariate fitted in the linear model. We also observed similar findings for the conditional GWAS analysis on chromosome 20 (S5 Fig).

By conducting a conditional analysis of slope traits on the intercepts, we detected multiple additional QTL signals (lead SNPs) across the genome at p < 1E-05 (S6 Fig). However, most of these lead SNPs were associated with a large FDR > 0.10 – FDR for each SNP computed following [27]. Of the few candidate variants (all of which were detected from HTFYslope traits) with FDR < 0.10, the strongest GWAS signal was in BTA 14 ~1.7 Mb, of which the lead SNP (Chr14:1726184) mapped to the downstream region of JRK (Jrk helix-turn-helix protein). Notably, this gene was found to regulate behavioural rhythms in *Drosophila* flies, which is crucial for adaptive response to environmental changes such as temperature variations [28].

When combining conditional GWAS results for slope traits (conditioning on the intercept traits) in the meta-analysis approach, we detected 40 lead SNPs (p < 1E-05), all of which associated with low FDR < 0.10 (S7 Fig and S3 Table). The mean LD between these 40 lead SNPs and the lead SNPs detected for intercept traits was very low (r^2^ < 0.20), confirming that the conditional analysis was successful in identifying additional candidate variants for heat tolerance (besides the QTL detected from the first-round of GWAS) that are not strongly associated with the level of milk production. The most significant lead SNP (Chr14:531267; p = 9.04E-12) mapped to the upstream region of the SLC39A4 gene, a member of the solute carrier family, required for intestinal zinc uptake.

### Candidate causal variants for heat tolerance across all analyses

The candidate causal variants for heat tolerance were defined as the lead SNP (most significant SNP within 5 Mb QTL window) plus other significant SNPs in strong LD (r^2^ > 0.8) with the lead SNP, 500 kb up or downstream of the chromosome. We identified a total of 3,010 candidate causal variants for heat tolerance (slope traits) across all the analyses: single-trait GWAS; a meta-analysis of single-trait GWAS results; and meta-analysis of conditional GWAS results for slope traits, most of which were intergenic (N = 1545; 51%) followed by intronic (N = 947; 32%) and upstream (N = 277, 9%) variants (Fig 3 and S1 Table). At least 25 candidate SNPs were missense variants, most (N = 13) of which were in chromosome 14, including two variants (Chr14:615597 and Chr14:616087) mapping to HSF1 (heat shock factor 1) gene.

**Fig 3.**
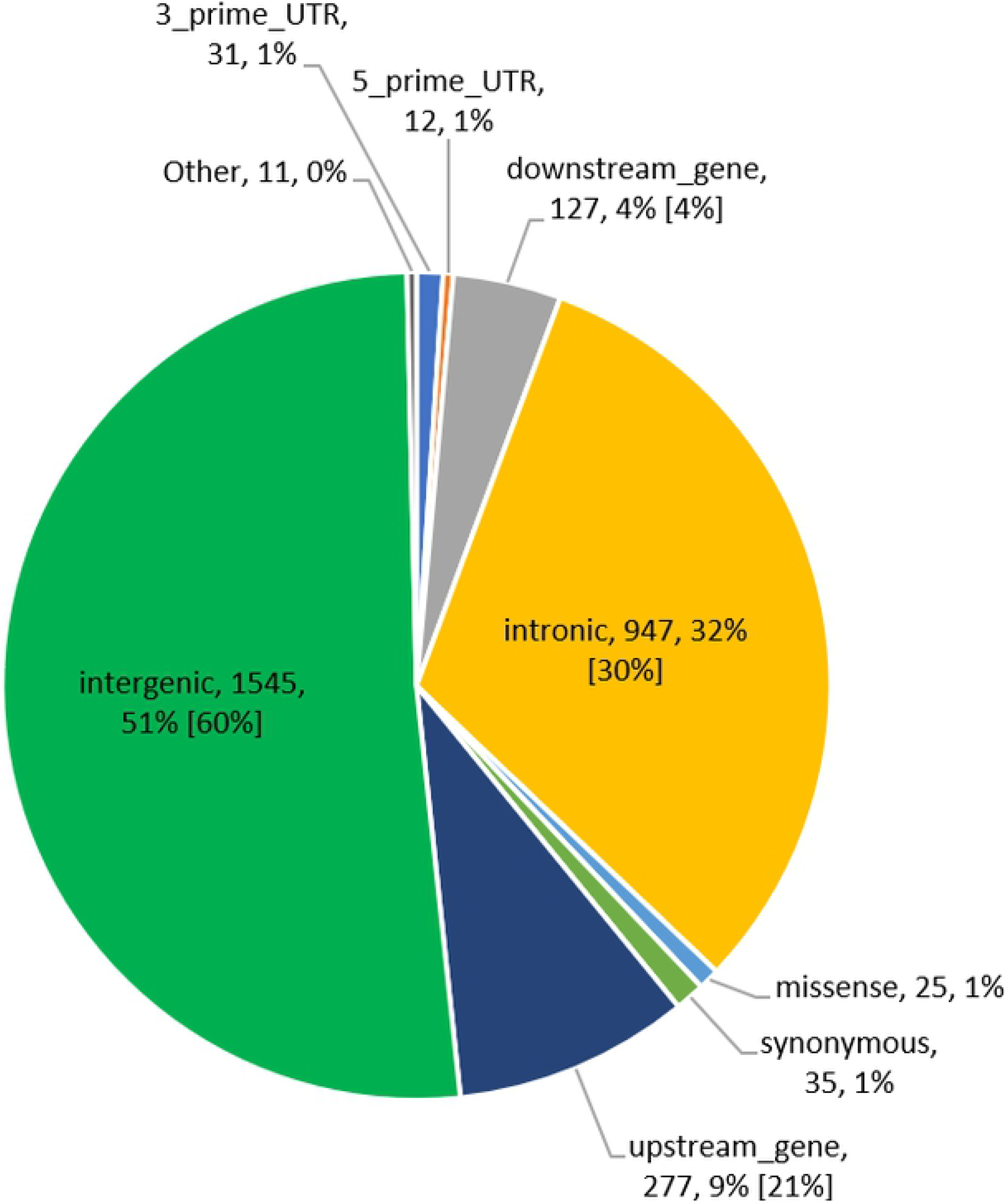
Proportion of candidate causal variants for heat tolerance within different functional classes identified from a) single-trait GWAS, b) meta-analysis, and c) meta-analysis of conditional GWAS results for slope traits. Values in brackets are the proportions of all variants used in the study (~15 million SNPs). Functional classes without values in brackets were represented by a small (< 1%) proportion of SNPs in the study dataset.

The candidate causal variants for heat tolerance are highly enriched (p = 8.54E-25) in the upstream gene regions (Fig 4), which agrees with GWAS for quantitative traits in humans [29], suggesting that they perhaps play a functional role in regulating gene expression. As expected, most candidate variants have modifier SnpEff [30] predicted impact (S5 Table). Two candidate causal mutations detected from the meta-analysis of slope traits have a high SnpEff predicted impact: a) a stop-gain mutation (Chr5:31184185) causing a premature stop codon in the *LALBA* (lactalbumin alpha) gene and b) a frameshift mutation (Chr29:41139622) in *STX5* (syntaxin-5) gene. The two candidate mutations appear to have a stronger effect on milk production compared to heat tolerance. This is evidenced by a smaller (p = 1.39E-19) p-value for the stop-gain mutation (Chr5:31184185) observed in the meta-analysis of intercept traits compared to the meta-analysis of slope traits (p = 4.08-12). Similarly, the p-value for the frameshift mutation (Chr29:41139622) in the *STX5* gene was smaller (p = 2.06E-16) for the meta-analysis of intercept traits than the meta-analysis of slope traits (p = 5.06E-06). None of these two candidate stop-gain mutations were significant (p < 0.05) following conditional GWAS for slope traits on intercept traits.

**Fig 4.**
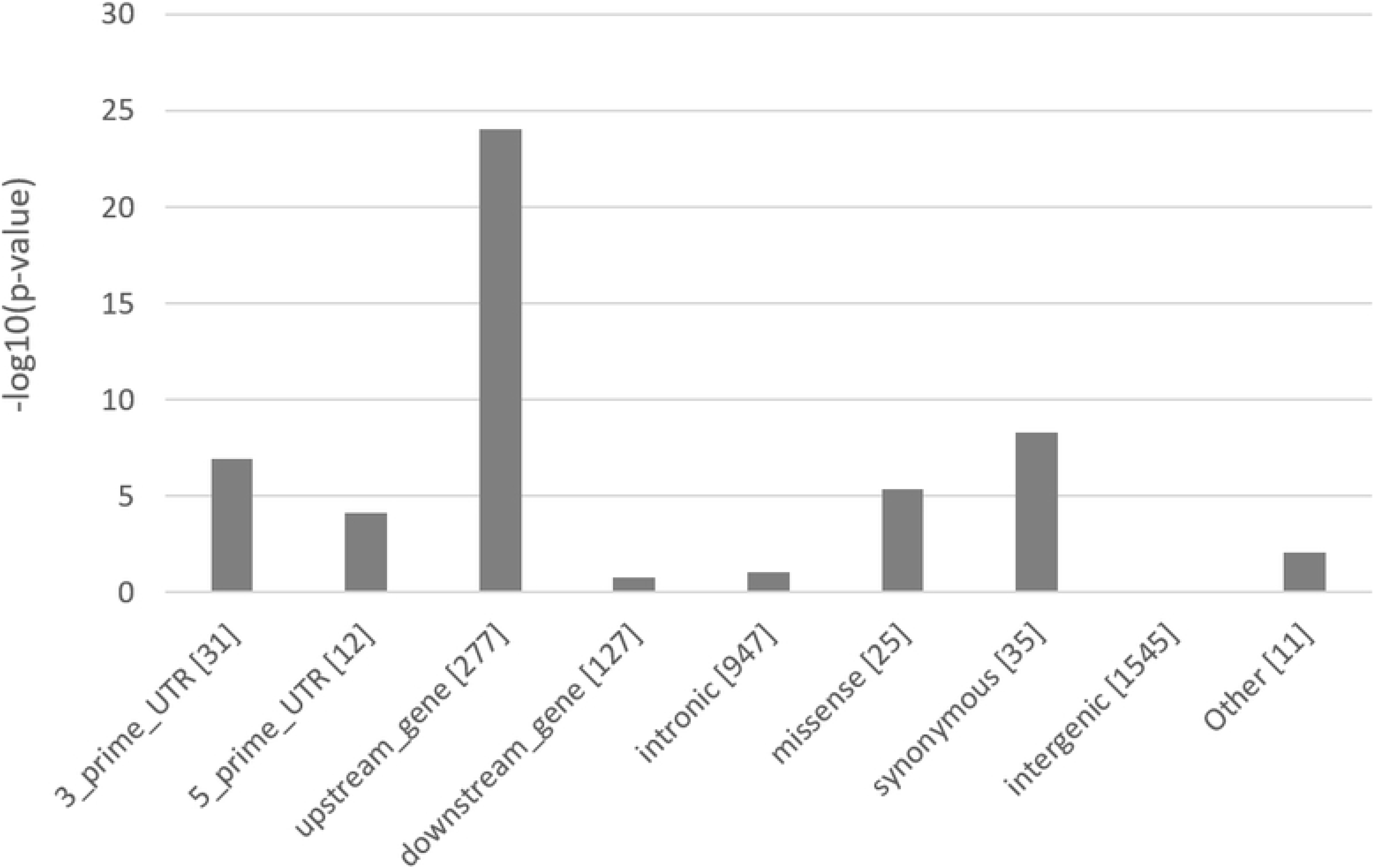
Enrichment of the candidate causal variants for heat tolerance across functional classes. The values in brackets are the number of variants within each class. The class “Other” includes variants with very small proportions of candidate variants (frameshift, stop-codon, splice variants, etc.).

Using data from [31], which documented over 300k sequence variants in cattle at highly evolutionarily conserved genome regions across 100 vertebrates (conservation/PhastCon scores > 0.9; see methods), we identified 61 potential functional variants for heat tolerance at these conserved sites in our study (S4 Table). However, the candidate causal mutations for heat tolerance are not enriched (p = 1.0) in the conserved regions of the genome.

Table 3 provides a short list of putative causal variants (upstream and missense) for heat tolerance that overlap at genomic sites highly conserved across vertebrates. Some of the candidate genes flanking these variants have been reported to be involved with cell survival under stress in animals, e.g., *SCD* [32], *KIAA1324* [33], and *TONSL* [15]. The *SCD* (stearoyl-CoA desaturase) gene encode fatty acid metabolic enzyme and perhaps is required for metabolic homeostasis during heat stress in mammals. Other putative candidate genes for heat tolerance include *KIFC2*, *VPS13B*, and *USP3*. For example, [34] demonstrated that the *USP3* gene, a member of the ubiquitin-specific proteases (USPs) family, is required for eliminating misfolded proteins under heat stress conditions in Yeast.

**Table 3.** Upstream and missense candidate causal variants for heat tolerance (slope) traits at genomic sites that are highly conserved (conservation score > 0.9) across 100 vertebrate species*.

| SNP | Chr | BP | Annotation | VEP_impact | Gene | HTMYslope | HTFYslope | HTPYslope | Meta-analysis<sup>1</sup> | Conditional analysis<sup>2</sup> | p-value<sup>3</sup> |
| --- | --- | --- | --- | --- | --- | --- | --- | --- | --- | --- | --- |
| rs209684414 | 3 | 34215670 | upstream | MODIFIER | *KIAA1324* |  |  |  | ✓ |  | 5.98E-07 |
| rs207668220 | 3 | 34273899 | upstream | MODIFIER | *C3H1orf194* |  |  |  | ✓ |  | 6.5E-07 |
| rs210324395 | 10 | 46503113 | upstream | MODIFIER | *USP3* |  |  |  | ✓ |  | 8.66E-09 |
| rs210468775 | 10 | 46505212 | upstream | MODIFIER | *USP3* |  |  |  | ✓ |  | 1.89E-09 |
| rs207681599 | 14 | 432274 | upstream | MODIFIER | *LRRC14* | ✓ | ✓ | ✓ | ✓ | ✓ | 1.46E-52\|1.13E-63\|1.13E-28\|8.8E-243\|7.92E-08 |
| rs136474298 | 14 | 471951 | missense | MODERATE | *KIFC2* | ✓ | ✓ | ✓ | ✓ | ✓ | 6.11E-53\|4.83E-63\|5.59E-29\|2.5E-242\|7.77E-08 |
| rs207886320 | 14 | 479761 | upstream | MODIFIER | *KIFC2* | ✓ | ✓ | ✓ | ✓ | ✓ | 6.18E-52\|4.17E-64\|4.55E-28\|3.9E-242\|7.85E-08 |
| rs137472016 | 14 | 494621 | upstream | MODIFIER | *TONSL* | ✓ | ✓ | ✓ | ✓ | ✓ | 6.28E-53\|8.39E-63\|5.94E-29\|9.1E-242\|7.87E-08 |
| rs445616049 | 14 | 64454721 | missense | MODERATE | *VPS13B* | ✓ |  |  | ✓ |  | 4.18E-08\|1.75E-32 |
| rs41946451 | 20 | 37085370 | missense | MODERATE | *CPLANE1* |  |  |  | ✓ | ✓ | 1.14E-07 |
| rs41255693 | 26 | 21272422 | missense&splice | MODERATE | *SCD* |  | ✓ |  | ✓ |  | 4.11E-07\|1.85E-08 |

\*Conservation scores (PhastCon score) of variants at conserved genomic sites were computed by [31] based on the LiftOver (https://genome.ucsc.edu/cgi-bin/hgLiftOver) human sites calculated across 100 vertebrate species; single-trait GWAS for heat tolerance milk (**HTMYslope**), fat (**HTFYslope**), and protein (**HTPYslope**) yield slope traits; <sup>1</sup>Meta-analysis combining single-trait GWAS results for slope traits; <sup>2</sup>Meta-analysis combining single-trait conditional GWAS results for slope traits; <sup>3</sup>For each analysis that identified this variant as significant.

### Pathway enrichment analysis

We generated a list of candidate genes mapping within or near lead SNPs detected at FDR < 0.10 for each trait set for the pathway enrichment analyses. We found that the candidate gene-list for slope traits were highly enriched for the KEGG pathways related to the neuronal system (neuroactive ligand-receptor interaction and glutamatergic synapse) and metabolism system (citrate cycle) (Fig 6). Interestingly, the heat tolerance candidate gene-list (N = ~ 400 genes) identified from various analyses (single-trait GWAS, meta-analysis, and conditional analysis) were consistently significantly enriched for a neuroactive ligand-receptor interaction pathway comprising of 15 genes (*CALCR, PTGER2, THRB, GRIK2, NPY2R, F2RL1, GRIN2A, NR3C1, CHRM3, GRM8, GRM7, GRID2, NPFFR2, MC4R, GHR*). A total of 8 genes were enriched (p = 4.0E-03) in the glutamatergic synapse pathway (*GRIN2A, GRM7, GRM8, ITPR1, ITPR2, SLC17A6, GRIK2, GRIA4*). The citrate cycle pathway was also enriched (p = 1.87E-03), comprising of 5 candidate genes for heat tolerance (*ACLY, PDHA2, MDH1, SUCLG2, PCK1*).

We also analysed a smaller set of genes (N = ~230) with the strongest (p < 1E-05) evidence of association for heat tolerance, separately (that is, the gene-list underlying the candidate causal variants defined as the lead SNP (most significant) within an independent QTL plus other significant SNPs in strong LD (r^2^ > 0.80) with the lead SNP, 500 kb up or downstream), to see enriched biological pathways. Interestingly, we observed enrichment (p = 0.02) of the genes in the neuroactive ligand-receptor interaction pathway, which provides strong support that this neuronal pathway is relevant for heat tolerance comprising of 8 genes (*GHR, NPFFR2, P2RY8, GRIN2A, CHRM1, THRB, CALCR, F2RL1*).

When examining the candidate gene-list from single-trait GWAS analyses for slope traits separately, the neuroactive ligand-receptor interaction pathway was overrepresented for candidate gene-list for HTMYslope (p = 3.19E-04) and HTPYslope (p = 7.79E-03) traits (Fig 7). On the other hand, gene-list for HTFYslope were enriched (p = 1.55E-02) for the axon guidance pathway comprising four genes (*ABLIM2, ABLIM3, NTN1, ROBO1*) and metabolic (p = 0.06) pathways.

To further test whether the neuronal pathway is real and not an artifact of our analyses for heat tolerance traits (slopes), we performed enrichment analyses for the significant candidate gene-list for intercepts traits (level of milk production traits). In the candidate gene-list for intercept traits, we found no evidence for enrichment (p < 0.05) in any neuronal pathways; thus, providing further favourable support that neuronal pathways are relevant for heat tolerance in mammals.

## Discussion

In this study, we performed a GWAS using a large sample size of Australian dairy cows (N = 29,107) with milk production records and imputed sequence data (~15 million SNPs) to identify candidate causal variants and functional genes and pathways associated with heat tolerance. Australia’s dairy cattle are uniquely placed for studying heat tolerance in mammals for two main reasons: 1) they are subjected to a wide range of seasonal climatic variations across diverse dairying regions spanning one of the geographically largest countries in the world, and 2) Australia’s dairying is predominantly pasture-based with limited heat stress mitigation measures in contrast with those, for example, in North America, where extensive managerial strategies are used more to reduce thermal stress. Overall, we have identified novel candidate causal variants in the neuronal pathways that contribute significantly to heat tolerance in animals.

We leveraged two statistical approaches to identify genetic loci and pathways for heat tolerance: single-trait GWAS linear models and multi-trait meta-analysis. Single-trait GWAS is based on regressing phenotypes on each SNP one at a time. On the other hand, a meta-analysis that combines results of the single-trait GWAS allowed us to discern putative pleiotropic genetic variants for heat tolerance. Consequently, we identified multiple novel loci for heat tolerance, including 61 potential functional variants at genomic sites highly conserved across 100 vertebrates (Table 3 and S4 Table), which could be valuable for fine-mapping and genomic prediction. Studies in humans [35] and cattle [31] have demonstrated that the conserved genomic sites have strong enrichment of trait heritability. Moreover, the results revealed specific candidate causal variants and genes related to neuronal functions for heat tolerance in animals, which we now discuss in more detail.

Heat stress responses are complex adaptations in animals involving many biological pathways, including the nervous system, which connects the internal and external environment to maintain stable core body temperature [36]. Among the candidate gene-list that contribute significantly to heat tolerance in the study animals (Holstein cows), the neuroactive ligand-receptor interaction and glutamatergic synapse pathways (Fig 6), as components of the nervous system, were highly enriched (p < 1E-03) biological features.

At least two candidate variants in the intronic region of *ITPR2* (Chr5:83330185; p = 1.3E-05) and *GRIA4* (Chr15:2461074; p = 5.8E-05) genes in the glutamatergic synapse pathway could be potential targets for resilience to environmental stress in animals. *ITPR2* gene was associated with heat stress in the US Holsteins [15] or sweating rate in humans and mice [37], while the *GRIA4* gene has been linked to thermoregulation in the Siberian cattle [38]. Another candidate variant (Chr22:21783956) detected for heat tolerance milk (p = 3.87E-05) and protein (7.15E-05) yield slope traits mapped to the intronic region of *ITPR1* – a gene associated with environmental adaptation in the domestic yak [39]. These three lead SNPs for slope traits overlapped with those for intercept traits, with opposing effect direction, suggesting that they affect both milk production and heat tolerance traits.

Previous studies show that the neuroactive ligand-receptor interaction is involved in maintaining energy homeostasis during heat stress in ducks [40]. As protein production is the most valuable output from dairy farms, the focus of breeding programs has been traits associated with yield, with the average milk volume per cow/year almost doubling within the past three decades in Australia [41]. The environmental heat stress, coupled with the elevated metabolic-induced thermogenesis, means that the genetic and cellular reprogramming of pathways such as the nervous system may be necessary to regulate a cascade of hormonal processes such as growth factors, insulin, serotonin, thyroid, prolactin, and mineralocorticoids associated with milk synthesis [42]. We identified 15 genes (FDR < 0.10) associated with the neuroactive ligand-receptor interaction, which could be relevant for metabolic homeostasis in cattle during thermal stress, of which three candidate genes (*GHR, NPFFR2,* and *CALCR*) showed the strongest evidence (p < 1E-05).

Here we discuss the evidence for each of these three candidate genes:

1) [21] demonstrated that the *NPFFR2* (neuropeptide FF receptor-2) gene, which is mainly expressed by neurons in the brain, plays a crucial role in regulating diet-induced thermogenesis and bone homeostasis in mice. In this study, two lead SNPs (Chr6:87070486 and Chr6:87249592), detected from single-trait GWAS for HTMYslope and HTPYslope (p < 1E-05) mapped to the intergenic and intronic regions of *NPFFR2* gene in BTA 6, respectively. Physiological studies suggest that NPFF family genes regulate feeding behaviour and energy expenditure in mammals [reviewed in 43]. During heat events, dairy cattle typically reduce their dry matter intake by up to 30%, perhaps as part of an adaptive mechanism to depress metabolic heat production [44]. Other studies, e.g., [45] show that inhibition of NPFF receptors induces hypothermia in mice. A recent review [46] indicates that NPFF and its receptors have many promising therapeutic applications including pain, cardiovascular, and feeding regulations in mammals. By examining the genomic region around the *NPFFR2* gene (Fig 5), it is more likely that the two lead SNPs within this QTL represent separate candidate causal mutations since they are not in strong LD. Interestingly, although the lead SNP (Chr6:87070486) for slope trait overlapped with the lead SNP detected for the milk yield intercept (MYint), we observed stronger evidence for the slope (HTMYslope; p = 3.05E-13) than the intercept (MYint; p = 4.19E-10), suggesting that this SNP is a good candidate for heat tolerance. Besides, this lead SNP (Chr6:87070486) remained significant (p = 6.36E-06) following single-trait conditional GWAS analysis for HTMYslope trait (conditioning slopes on the intercept traits) as well as in the meta-analysis of single-trait conditional GWAS results for slope traits (p = 3.74E-06).
2) Calcitonin receptors regulate daily body temperature rhythm in mammals and insects and are essential for maintaining homeostasis [47]. In this study, the lead SNP (Chr4:10815768) was intronic in the *CALCR* (calcitonin receptor) gene, perhaps indicating that it could be relevant for animals experiencing recurrent or chronic stress, such as in Australian seasonal summers. The strong GWAS signal around this QTL (S8 Fig) suggests that the *CALCR* gene likely harbours causal mutations affecting heat tolerance. Dairy cattle employ various adaptive behavioural strategies during heat stress such as reduced feed intake, increased volume, and frequency of water intake, increased standing time, shade seeking, and grazing at cooler day time. We think that *CALCR* is likely involved with some of these heat-stress adaptive behaviours in dairy cattle. Future studies are needed to confirm this, particularly by combining production traits with other relevant behavioural phenotypes such as panting scores from high-throughput recording devices, e.g., activity-based collars.
3) The expression of the *GHR* (growth hormone receptor) gene is down-regulated during heat stress in livestock, including dairy cows [48] and avian species [40]. The adaptive physiological significance of this down-regulation is not well understood, and it is partly independent of the nutritional level of the animal [48]. In this study, the lead SNP (Chr20:32103408; p = 2.01E-08) identified only in one slope trait (HTMYslope) based on significant cut-off of p < 1E-05 mapped to intronic region of *GHR* gene (S9 Fig). However, we found a stronger signal after combining the GWAS results for all the slope traits in a meta-analysis with the lead SNP (Chr20:32201287; p = 1.7E-47) mapping to the intergenic (~22 kb) region of the *GHR* gene, which confirms the pleiotropic effect of this QTL [49]. Also, we observed no significant SNP (p < 1E-05) around this QTL following single-trait conditional analyses, but a somewhat strong signal emerged when we combined single-trait conditional GWAS results in the meta-analysis, for which the lead SNP (Chr20:32226298; p = 5.35E-07) mapped to the intergenic region (~47 kb) of GHR. This further supports a possible second QTL that is independent of the level of milk production and shows pleiotropy for the heat tolerance traits. Other published GWAS have also reported an association of the GHR gene with milk production in heat-stressed cows [15] and respiratory rates in pigs during heat stress [17]. Several studies have also implicated the *GHR* polymorphisms to milk production in bovines, e.g., Chr20:31888449 phenylalanine-to-tyrosine missense mutation [26]. This mutation was not in strong LD (r^2^ > 0.8) with the lead SNP detected for slope traits in our study. Taken together, polymorphisms around the *GHR* gene could be candidate targets for improving thermotolerance in livestock, although with possible antagonistic effect on milk production considering, for example, the opposing effect direction observed for the lead SNP (Chr20:32103408) within this QTL on the slope (HTMYslope) and intercept (MYint) traits.

**Fig 5.**
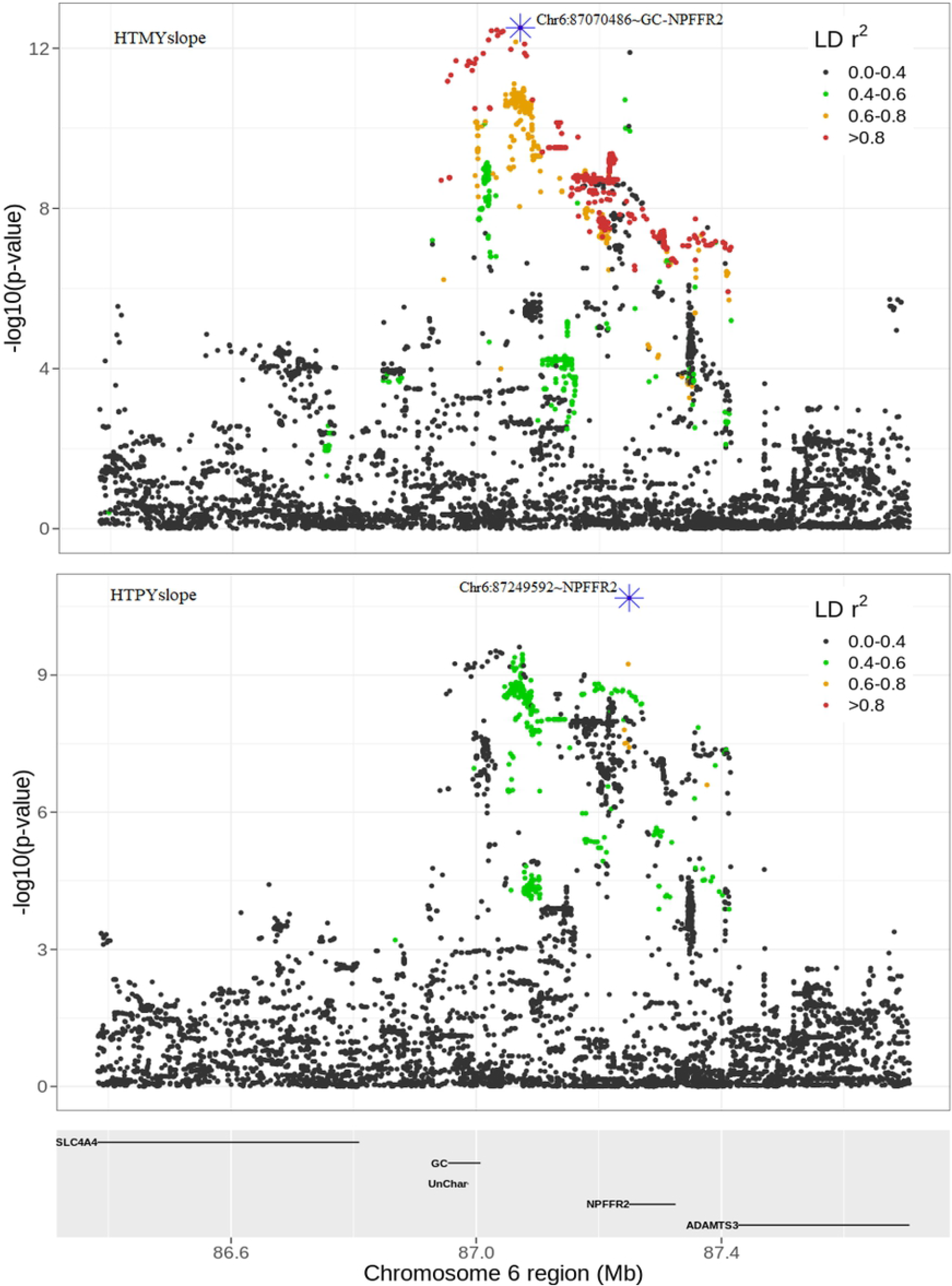
QTL discovery for heat tolerance milk (HTMYslope) and protein (HTPYslope) yield slope traits around the *NPFFR2* gene in bovine chromosome 6.

There is general agreement that heat stress decreases milk yields (milk, proteins, fat, etc.) in dairy cattle. However, the genetic and biological basis for this reduction is still unclear. Evidence suggests that the reduced feed intake in heat-stressed dairy cows is independent of reduced milk yields and composition [44]. The molecular control and pathways for individual milk traits during heat stress are scarce and inconclusive. In this study, the QTLs detected for the heat tolerance traits varied across the three milk traits (HTMYslope, HTFYslope, HTPYslope), suggesting that they are, in part, regulated by different genes in heat-stressed cows. The greater overlap of candidate genes observed for HTMYslope and HTPYslope traits was expected due to their relatively high correlation (0.90) compared to HTMYslope and HTFYslope (0.56) or HTPYslope and HTFYslope (0.62). These correlations appear to mirror the proportions of SNPs with the same or inconsistent effect direction observed for significant SNPs between slope traits. Considering that heat stress alters carbohydrate, lipid, and amino acid metabolism [50], the large proportion of SNPs with inconsistent effect direction, particularly between HTPYslope and HTFYslope, suggest that these traits are somewhat differently regulated in heat-stressed dairy cows.

Several pair-fed studies suggest that pathways related to the mammary gland protein synthesis govern protein production under heat stress in dairy cows, in part, via reduced amino acid supply to the mammary gland, e.g., [51, 52]. We found that the candidate genes for HTMYslope and HTPYslope traits were overrepresented (p < 0.005) in the neuroactive ligand-receptor interaction pathway. This agrees with [53] that genes associated with milk proteins are involved in neuronal signaling pathways in dairy cattle. However, it remains unclear how this pathway is regulated during heat stress conditions in dairy cows to impact protein production.

On the other hand, the molecular pathways for fat production under heat stress conditions have not been widely studied. Some studies [e.g., 54] suggest that the reduced activation of PPAR (peroxisome proliferator-activated receptor) signaling pathways leads to decreased expression of genes associated with fat metabolism. Candidate genes for HTFYslope identified in this study are associated with the KEGG term “metabolic pathways” (Fig 7). Five candidate genes (*DMGDH, PDHA2, UGP2, MDH1, PRDX6, NDUFA13*) within this pathway may be involved with alleviating oxidative stress in heat-stressed cows. In line with these findings, we found that the candidate genes for heat tolerance (Fig 6) are overrepresented in the citrate cycle/TCA pathway, which is central to mitochondrion energetics, and might serve to reduce substrate oxidation and reactive oxygen species (ROS) production, thus preventing cellular damage during heat stress.

**Fig 6.**
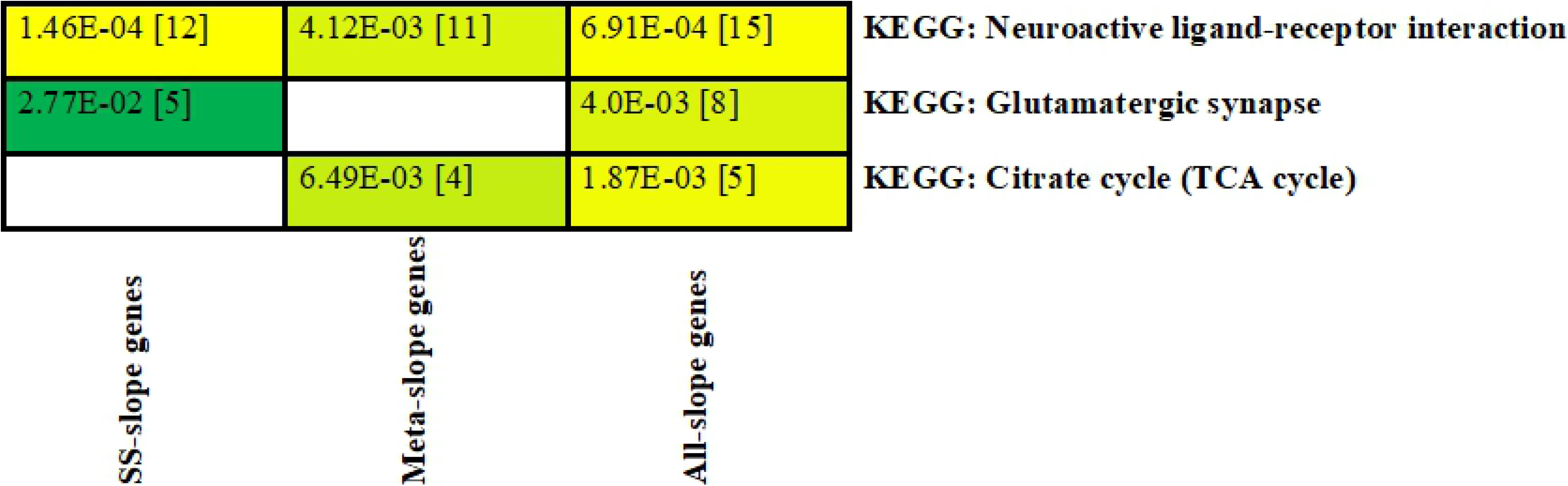
Enriched Kyoto Encyclopedia of Gene and Genomes (KEGG) pathways obtained from candidate gene-list for slope traits detected at false discovery rate (FDR < 0.10): SS-slope genes – gene-list from single-trait GWAS; Meta-slope genes – gene-list from multi-trait meta-analysis of slope traits; All-slope genes – combined gene-list from single-trait and meta-analysis. Cells are color-coded according to the strength of the significance for each pathway. Values in brackets are the number of genes within each pathway.

**Fig 7.**
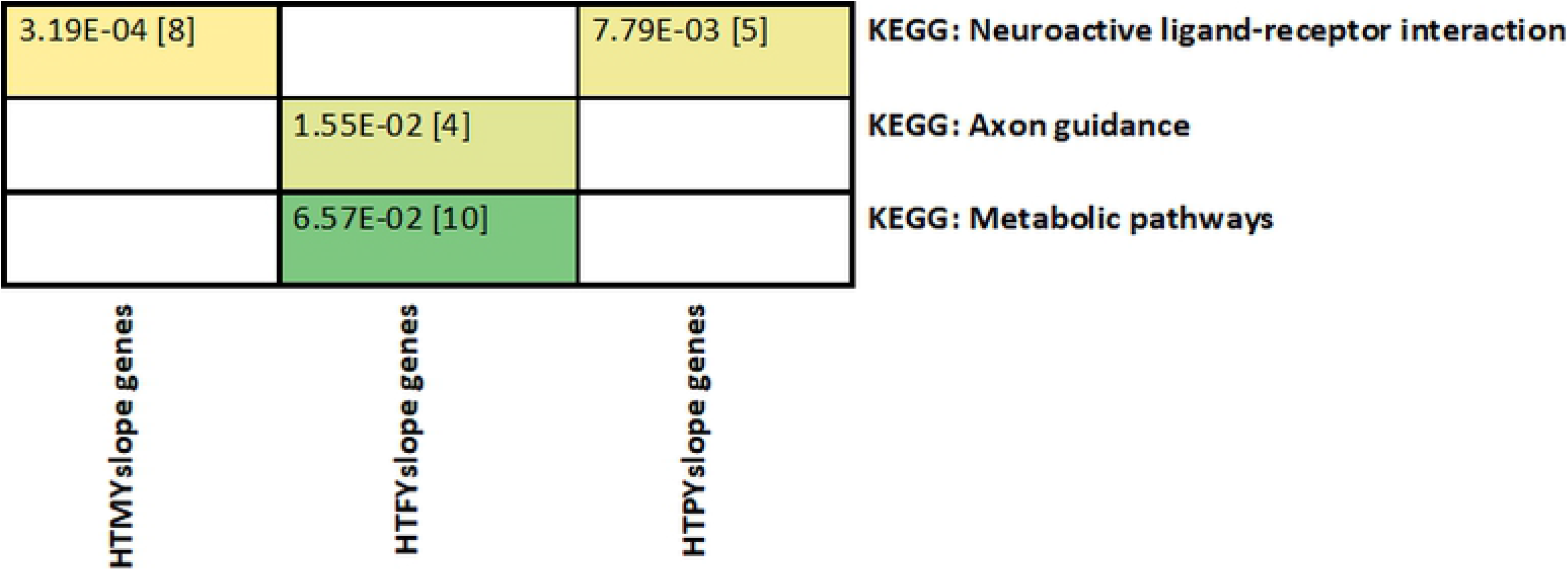
Enriched Kyoto Encyclopedia of Gene and Genomes (KEGG) pathways obtained from our gene-list for single-trait GWAS analysis of slope traits: HTMYslope (heat tolerance milk yield slope); HTFYslope (heat tolerance fat yield slope); and HTPYslope (heat tolerance protein yield slope). Cells are color-coded according to the strength of the significance for each pathway. Values in brackets are the number of genes within each pathway.

Notably, our pathway results are perhaps not directly comparable to most previous work in which the study cows were subjected to short-term acute heat stress under experimental conditions [e.g., 54] whereas the current work mimics recurrent or chronic stress that dairy cows experience during summer seasons in Australia. The effects of heat stress in livestock depend on its duration and severity, with the most recent work in Arabian camels somatic cells showing that acute heat stress elevates the expression of heat shock proteins and DNA repair enzymes while chronic heat leads to changes in cell integrity and reduction of total protein levels, metabolic enzymes, and cytoskeletal proteins [55]. Our candidate QTLs are particularly important since it provides novel insights into the molecular aspect of chronic stress considering that the study animals are predominantly reared under outdoor conditions with limited heat stress mitigations. Future studies are required to confirm if these QTLs are involved with recurrent chronic stress in other animal species.

We could not replicate most of the candidate genes with published GWAS results for heat tolerance in cattle, likely for several reasons. First, all comparable earlier studies were much smaller (< 5,000 animals) and therefore were under-powered, and the marker density used was typically 50k or 600k SNP array [e.g., 14, 15]. As expected, we observed that our sequence variants showed markedly higher significance levels than the 50k SNP array and increased the number of significant peaks across the genome (S1 Fig). Second, the trait used to define heat tolerance in this study (i.e., the rate of milk yield decline under heat stress) differs from many other studies [e.g., 13], which used measures of core body temperatures in their GWAS. Given that heat tolerance is a complex trait involving a wide array of adaptative responses (behavioural, physiological, cellular, etc.), different QTLs may be captured by different traits used in GWAS. Third, differences in the patterns of LD among study populations used and imputation quality may have implications on GWAS, particularly in the detection of putative causal mutations [56]. Here we explored QTLs for heat tolerance in purebred Holstein cows, while some other studies, e.g., [16] have used crossbred cattle. Collectively, these factors likely impacted the replication of previous GWAS candidate genes for heat tolerance.

Although we detected multiple candidate causal variants for heat tolerance in this study, it appears that larger sample size (we used N = 29,107) would be beneficial considering the polygenic architecture of this trait. Larger sample size is required to detect causal variants with very small effects and the effects of rare causal variants [12]. For example, many of the lead SNPs (most significant) for heat tolerance were tagged by none or very few significant SNPs (S1 Table), which may be false-positive variants passing the GWAS cut-off (p < 1E-05). Our results support the highly polygenic nature of heat tolerance characterised by multiple small-effect variants, suggesting that this trait is more amenable to genomic selection tools such as those currently implemented in the Australian dairy industry [3, 57] rather than approaches that exploit few QTLs with large effects.

In conclusion, we performed GWAS for heat tolerance using large sample size and genotype dataset for dairy cattle. The increased sample size and high-resolution SNP data in our study compared to previous reports allowed us unprecedented power and precision of the GWAS to pinpoint multiple putative causal mutations, including 61 potential functional variants at genomic sites highly conserved across 100 vertebrate species. Also, results indicate that different genes and pathways, in part, regulate different milk production traits (milk, fat, and proteins) in heat-stressed dairy cows with a substantial overlap of genes for heat tolerance milk and protein yields. Overall, the results revealed the importance of variation in genes related to the neuronal functions for heat tolerance in mammals, which is of interest for future research towards understanding and managing heat stress for warm climates and particularly in view of the anticipated climate changes.

## Materials and methods

### Animals and phenotypes

No live animals were used in this study. Phenotypes used for GWAS were part of our previous study [58] obtained from DataGene (DataGene Ltd., Melbourne, Australia; https://datagene.com.au/) – the organisation responsible for genetic evaluation of dairy animals in Australia. The phenotypes were test-day milk, fat, and protein yields for Holstein dairy cows collected from dairy herds that were matched with climate data (daily temperature and humidity) obtained from weather stations across Australia’s dairying regions. The distribution of dairy herds and weather stations; and the calculation of environmental covariate (i.e., temperature-humidity index (THI)) used here were described in our earlier studies [3, 58].

### Calculation of heat tolerance phenotypes for cows

The dataset used to calculate heat tolerance phenotypes for cows was similar to that used by [58], comprising a total of 424,846 test-day milk records for first, second and third lactations from 312 herds and 15,906 herd-test days (**HTD**) collected over 15 years (2003-2017). A summary of the final dataset is given in Table 1. The rate of decline (slope) in milk, fat, and protein yield due to heat stress events was estimated using a reaction norm models [58]. In these models, data on milk, fat, or protein yield were adjusted for the fixed effects, including herd test day, year season of calving, parity, age at calving, jointly for parity and DIM, and jointly for the stage of lactation and THI. Random effects fitted in the model included a random regression on a linear orthogonal polynomial of THI, where the intercept represents the level of mean milk yield and the linear component represents the change in milk yield (slope) due to heat stress for each cow and a residual term. In the model, the threshold of THI was set to 60 following [59]. The analyses to derive trait deviation (**TD**) which represents a phenotype adjusted for all fixed effects (i.e., the mean/intercept and slope for each cow) were conducted using ASReml v4.2 [60].

We refer to milk intercept traits as [MYint (i.e., milk yield intercept), FYint (i.e., fat yield intercept), and PYint (i.e., protein yield intercept)] and the slopes traits as [HTMYslope (i.e., heat tolerance milk yield slope), HTFYslope (i.e., heat tolerance fat yield slope), and HTPYslope (i.e., heat tolerance protein yield slope)], respectively.

### Genotypes

Two genotype datasets were analysed for 29,107 Holstein cows with the above phenotypes: 50k SNP chip and 15,098,486 imputed whole-genome sequence variants (**WGS**). Most of the cows were originally genotyped with a custom low-density 10k SNP panel or a standard medium density 50k SNP array (BovineSNP50k BeadChip: Illumina Inc). The low-density genotypes were imputed to the 50k array using a reference set of approximately 14,000 animals with real 50k genotypes, with approximately 7,000 SNPs of the low-density SNP panel overlapping the 50k SNP array. The 50k genotypes were then imputed to the high-density Bovine SNP array (HD: BovineHD BeadChip, Illumina Inc) using a reference set of 2,700 animals with real HD genotypes. All SNP BeadChip genotypes were first converted to the ARS-UDC1.2 reference genome (https://www.ncbi.nlm.nih.gov/assembly/GCF_002263795.1/) [61] positions from reference genome UMD3.1 and imputed using Fimpute3 [62]. The WGS was imputed from the HD genotypes using a reference set of 3,090 *Bos taurus* sequences in the Run7 of the 1000 Bull Genome Project (http://1000bullgenomes.com/) [18] aligned to the ARS-UCD1.2 reference genome. Only bi-allelic sequence variants with a minor allele count (≥ 4) and GATK [63] quality tranche 99.0 or better were retained for imputation. Pre-imputation, we also removed variants with higher than expected heterozygosity (> 0.5) if they fell in a 500 kb window enriched for variants showing excessive heterozygosity (as a proxy to indicate regions where WGS mapping/alignment may be poor). A total of 31,994,954 sequence variants remained for imputation. Minimac3 [64] was used for WGS imputation, having first pre-phased both the HD genotypes and the WGS reference using Eagle v2 [65]. For the analysis, we retained only the variants with Minimac3 imputation accuracy, *R*^2^ > 0.4 and MAF > 0.005 (N = 15,098,486 sequence variants).

### Single-trait GWAS and multi-trait meta-analysis

A genome-wide association analysis (**GWAS**) using a mixed linear model was used to test associations between individual SNP and cows’ slope [HTMYslope, HTPYslope and HTFYslope] and intercept [MYint, FYint, PYint] traits using GCTA software [66]. Because phenotypes were TD already adjusted for nongenetic effects, for each autosomal SNP *i* with minor allele frequency (**MAF**) > 0.005, the fitted model per trait was,

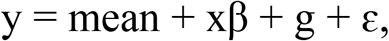

where **y** was the vector of TD for cows (n = 29,107), *β* was the allele substitution effect of SNP *i*, **x** was the vector of genotype dosages (0, 1, or 2) for SNP *i*, **g** was the vector of polygenic effect with 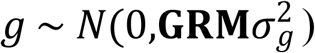 and ***ε*** was a vector of the residual effect with 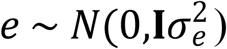, where **I** was an *n* × *n* identity matrix. The variance of **y** was 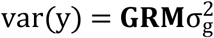 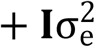 where **GRM** is the genomic relationship matrix between cows, and 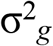 and 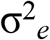 were the genetic and residual variances. For animal j and k relationship was calculated using GCTA [66] as follows:

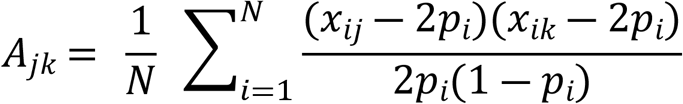

where *A*_*jk*_ are the off-diagonal elements of GRM for animal j and k; N = total number of SNPs from 50k SNP array data (MAF > 0.005; 45,504 SNPs); *x*_*ij*_ and *x*_*ik*_ are genotypes are the number of copies for reference allele for the *i*th SNP *j*th and *k*th cow; and *p*_*i*_ is the allele frequency for *i*th SNP.

Genomic heritability was calculated for each trait using variance component estimates from – reml option of GCTA for 50k SNP array (45,504 SNPs) data of cows (N = 29,107): 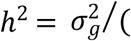 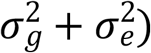.

To increase the power of GWAS and identify pleiotropic variants, we next combined single-trait GWAS results obtained above in a multi-trait meta-analysis following [19]. The multi-trait *χ*^2^ statistics for *i*th SNP was calculated separately for intercept [MYint, FYint, PYint] and slope [HTMYslope, HTFYslope, and HTPYslope] traits as follows:

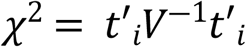

where *t*_*i*_ is the vector of 3 × 1 vector of signed t-values (i.e., b/se) of *i*th SNP for either intercept or slope traits; and *V*^−1^ is the inverse of 3 × 3 correlation matrix of the signed t-values calculated based on all pairs for the intercept or slope traits. The significance of *χ*^2^value for *i*th SNP was calculated based on chi-squared distribution with 3 degrees of freedom – that is number of traits for either intercept or slope traits.

### Conditional GWAS analysis

Next, we performed two conditional GWAS strategies of slope traits using GCTA software to test somewhat different hypotheses:

a) Conditional analysis of slope traits on lead SNP (i.e., most significant SNP within a chromosome from first-round GWAS) – aimed at identifying additional or secondary putative causal variants beside those detected from first-round GWAS. We performed a conditional analysis strategy on two chromosomes (BTA 14 and BTA 20), which showed the strongest GWAS signal for slope traits in the first-round GWAS (S1 and S2 Figs) and are known to harbour QTLs with major effects on milk production (i.e., BTA14 ~*DGAT1* and BTA20 ~*GHR* gene).
b) Conditional analysis of slope traits on intercept traits – aimed at identifying QTLs for heat tolerance that are independent, or not also strongly associated with the level of milk production. We fitted the intercept traits of MYint, FYint, and PYint, as a covariate in the linear model when analysing the HTMYslope, HTFYslope, or HTPYslope, respectively. To increase the power of GWAS, we then combined conditional GWAS results for the three slope traits [HTMYslope, HTFYslope, and HTPYslope] in a multi-trait meta-analysis following [19] as described earlier.

### Identifying candidate causal variants

We used the following criteria to select candidate variants (p < 1.0E-05) from the three analytical approaches (single-trait GWAS, meta-analysis, conditional analysis).

1. For each trait, select all SNPs with p < 1E-05.
2. Split each chromosome (N = 1…29) into 5 Mb non-overlapping windows from the start to the distal end of the chromosome.
3. Within the *i*th 5 Mb window, select the most significant SNP (i.e., the SNP with the smallest p-value below the threshold of p < 1E-05) defined as the ‘lead SNP’. We chose this arbitrary 5 Mb window size to obtain a small set of significant lead SNPs representing independent QTL (that is, not in linkage disequilibrium) for further detailed examination.
4. Calculate the LD between each lead SNP and all the other SNPs within 500 kb up and downstream of the lead SNP using Plink v1.9 [67].
5. For each lead SNP, extract all the significant SNPs (p < 1E-05) in strong LD (r^2^ > 0.80) with the lead SNP within 500 kb up or down downstream – to account for the fact that the lead SNP (most significant) is not necessarily the causal variant.

### Annotation of sequence variants and enrichment analysis

Annotation of all variants (~15 million SNPs) was performed using SnpEff [30] tool. Using the annotation, we grouped the candidate causal variants for heat tolerance (slopes) into 9 classes (intergenic, intronic, missense, upstream, downstream, 3_prime_UTR, synonymous, 5_prime_UTR, and Other) and performed enrichment analysis using phyper in R v3.61 [68]. The class “Other” comprised variants including 5_prime_UTR_premature/_start_codon_gain, frameshift, missense&splice, splice&intron, stop_gained, etc. S1 Table provides the number of candidate causal variants for heat tolerance within the 9 classes.

### Candidate variants at conserved genomic sites

We identified candidate causal variants for heat tolerance at highly conserved genomic sites using data from [31]. Briefly, these authors documented over 300k sequence variants at conserved sites in cattle based on the LiftOver (https://genome.ucsc.edu/cgi-bin/hgLiftOver) human sites with conservation scores (PhastCon score) > 0.9 calculated across 100 vertebrate species (see https://www.pnas.org/content/pnas/suppl/2019/09/07/1904159116.DCSupplemental/pnas.1904159116.sapp.pdf for more details).

### Pathway enrichment analysis

We generated candidate gene-list mapping near or underlying lead SNPs (most significant SNPs within 5 Mb QTL windows) identified at FDR < 0.10 cut-off threshold from both single-trait and multi-trait analyses of intercept or slope traits. For intergenic lead SNPs, we selected the closest gene on either side of the SNP. We chose this cut-off (FDR < 0.10) instead of a more stringent p < 1E-05 to include genes associated with smaller effects while guarding against false positives. We then performed the Kyoto Encyclopedia of Genes and Genomes (**KEGG**) enrichment analysis using DAVID [69].

We also performed enrichment test separately for the gene-list associated with potential major effects on heat tolerance identified across all analyses (i.e., gene-list with the strongest (p < 1E-05) evidence of association defined as the candidate causal variants (i.e., lead SNP + other significant SNPs in strong LD (r^2^ > 0.80) with the lead SNP within 500 kb up or downstream passing the cut-off p-value of 1 < 1E-05). For all the analyses, we considered functional pathways with Fisher’s p < 0.05 as significantly enriched.

## Acknowledgments

We are grateful to the 1000 Bull Genomes Project consortium for providing access to the Run7 cattle sequence data. We thank Bolormaa Sunduimijid (Agriculture Victoria Research) for the imputation of sequence data and Paul Stothard et al. from the University of Alberta for annotation of the sequence variants. Thanks to all farmers and DataGene Ltd. (Melbourne, Australia) for providing the phenotype data.

## Supporting information captions

**S1 Table. Number of candidate causal variants (p < 1E-05) for slope traits across different functional classes identified for** a) single-trait GWAS, b) meta-analysis of single-trait GWAS results, and c) meta-analysis of conditional single-trait GWAS results of slopes (conditioning each slope trait on the intercept traits).

**S2 Table. Candidate causal variants for heat tolerance identified from single-trait GWAS and multi-trait meta-analysis based on the GWAS cut-off threshold of p < 1E-05.**

**S3 Table. Candidate causal variants for heat tolerance detected following meta-analysis of conditional GWAS results for slope traits.**

**S4 Table. Candidate causal variants for heat tolerance at genomic sites highly conserved (conservation score > 0.9) across 100 vertebrate species.**

**S5 Table. Number of variants within different SnpEff predicted impact groups and enrichment scores.**

**S1 Fig. Manhattan plot of GWAS p-values for 29,107 Holstein cows based on 50k SNP set (left panel) and whole-genome sequence variants (right panel; N = 15,098,486 SNPs) for:** heat tolerance milk (HTMYslope; A), fat (HTFYslope; B) and protein (HTPYslope; C) yield slope traits. Dashed horizontal lines represent GWAS cut-off of p < 1E-05.

**S1 Fig. Manhattan plot of GWAS p-values for 29,107 Holstein cows obtained from 15 million imputed-WGS for** heat tolerance milk (HTMYslope; A), fat (HTFYslope; B), and protein (HTPYslope; C) yield slope traits. The highlighted red points are lead SNPs (most significant) identified using 5 Mb non-overlapping windows at p < 1E-05 (horizontal dashed line).

**S2 Fig. Overlap of candidate QTLs (p < 1E-05) from single-trait GWAS for heat tolerance milk (HTMYslope), fat (HTFYslope), and protein (HTPYslope) yield slope traits.** QTLs were defined as overlapping if the lead SNPs (most significant) within QTLs are close (within 1 Mb). The two QTLs which overlapped across the 3 slope traits are located around DGAT1 and MGST1 genes.

**S3 Fig. Distribution of linkage disequilibrium (LD) scores between lead SNPs (most significant) for slope traits and nearby (within 1 Mb) lead SNPs for intercept traits identified from single-trait GWAS and multi-trait meta-analyses.**

**S4 Fig. GWAS p-values on chromosome 20 at 30 to 36 Mb for heat tolerance milk yield slope (HTMYslope) trait.** The left plot (GWAS p-values before conditional analysis), middle (after conditioning slope on the lead SNP defined as the most significant SNP selected from first-round of GWAS; Chr20:32103408), and right plot (GWAS p-values after conditioning with milk yield intercept trait).

**S5 Fig. Conditional GWAS results for heat tolerance milk (HTMYslope), fat (HTFYslope), and protein (HTPYslope) yield slope traits.** GWAS for A, B and C were conditioned on milk, fat, and protein yield intercept traits, respectively. The highlighted red points represent the lead SNPs (most significant) within 5 Mb non-overlapping windows across the chromosome.

**S6 Fig. Manhattan plot of p-values obtained from combining conditional single-trait GWAS results for slope traits in the multi-trait meta-analysis.** The dashed line is the significant GWAS cut-off at p < 1E-05, while the red circles are the lead SNPs (most significant per QTL).

**S7 Fig. QTL discovery for heat tolerance protein yield slope (HTPYslope) trait around the CALCR gene region in bovine chromosome 4.**

**S8 Fig. QTL discovery for heat tolerance protein yield slope (HTPYslope) trait and meta-analysis of slope traits (Meta-HTslope) around the GHR gene region in bovine chromosome 20.**

## Data availability

Positions and annotations for all the lead SNPs (most significant SNPs) with p <1E-5 are in S1-S3 Tables. The study used third-party data obtained from DataGene (DataGene Ltd., Melbourne, Australia; https://datagene.com.au/). As strict agreements are in place between farmers and DataGene, this data is not publicly available. However, research related requests for access to the data may be accommodated on a case-by-case basis.

## Financial disclosure statement

This study was supported by DairyBio (Melbourne, Australia), funded by Dairy Australia (Melbourne, Australia), the Gardiner Foundation (Melbourne, Australia), and Agriculture Victoria (Melbourne, Australia). The funders had no role in study design, data collection and analysis, decision to publish, or preparation of the manuscript.

## Author contributions

JEP, HM, IMM, and EKC, conceived and designed the study; IMM assisted with preparation and imputation of genotype data; EKC, IMM, MH, JEP, BGC, RX, contributed to the formal data analysis; EKC wrote the first draft; all authors, reviewed and approved the final manuscript for publication.

## Competing interests

The authors declare no competing interests.

